# Human enteric glia diversity in health and disease: new avenues for the treatment of Hirschsprung disease

**DOI:** 10.1101/2023.09.26.559481

**Authors:** J.D. Windster, L.E. Kuil, N.J.M. Kakiailatu, A. Antanaviciute, A. Sacchetti, K. C. MacKenzie, J. Peulen-Zink, Tsung Wai Kan, E. Bindels, E. de Pater, M. Doukas, S. Yousefi, T.S. Barakat, C. Meeussen, C.E.J. Sloots, R.M.H. Wijnen, K. Parikh, W. Boesmans, V. Melotte, R.M.W. Hofstra, A. Simmons, M.M. Alves

## Abstract

Hirschsprung disease (HSCR) is caused by an absence of the enteric nervous system (ENS), which is crucial for intestinal function. The ENS is composed of enteric neurons and glia, and is mostly derived from migrating vagal neural crest cells. Trunk-derived Schwann cells also play a significant role in postnatal maintenance of the ENS. However, the diversity of the ENS in health and disease remains largely unknown. Here, we performed single cell RNA sequencing on pediatric controls and HSCR individuals, and identified two major classes of enteric glia, being canonical and Schwann-like enteric glia. We show that the latter are the main contributors of enteric glia heterogeneity after birth and importantly, that they are preserved in aganglionic segments of HSCR individuals. In a zebrafish model of HSCR, which also shows preservation of Schwann-like enteric glia, enteric neurogenesis could be stimulated, demonstrating a potential novel therapy for HSCR.

## Introduction

The enteric nervous system (ENS) is a major part of the autonomic nervous system that is responsible for the intrinsic innervation of the gastrointestinal (GI) tract. It consists of interconnected networks (plexuses) of enteric neurons and enteric glia cells that extend along the entire length of the intestine (1), and has a major role in regulating GI function and homeostasis. The ENS is for the most part, formed by one rostro-caudal wave of migrating neural crest-derived precursors (NCDPs) that originate from the vagal neural crest (2). After colonization by NCDPs, extrinsic nerve fibers from both cranial and trunk regions, reach the GI tract and Schwann cell precursors (SCPs) enter the gut (3, 4). SCPs are a neural crest-derived stem cell pool found in the peripheral nervous system, which contribute to different cell types, such as melanocytes, sympathetic and parasympathetic neurons, neuroendocrine cells (5), and ENS cells (6, 7). Interestingly, SCPs have been shown to be a major contributor of enteric neurogenesis postnatally, contributing to 5% of the submucosal neurons in the small intestine and around 20% of neurons in the colon of adult mice (7). In humans, enteric glia are expected to be up to seven times more abundant than neurons (8), and depending on the intestinal microenvironment, they present distinct morphological features. To date, up to six main types of enteric glia have been identified in humans, based on their location (intra or extraganglionic) and morphology (9).

Defects in the early establishment of the ENS due to abnormal proliferation, survival, migration or differentiation of enteric neural crest cells, are associated with numerous congenital intestinal neuropathies, including Hirschsprung disease (HSCR). HSCR is a complex genetic disorder, with an incidence of 1:4500 live births (10). To date, pathogenic variants in more than 20 genes have been identified to contribute to the genetic etiology of this disease (11). However, the Rearranged during transfection (*RET*) gene takes center stage, as it accounts for the majority of cases (50 % of familial and 15-20% of sporadic cases) (12), and is associated with the most severe forms of HSCR (total colonic or total intestinal aganglionosis) (13). Interestingly, although the aganglionic segment of HSCR individuals is devoid of enteric ganglia, a characteristic feature of HSCR disease is the presence of extrinsic-derived neural fibers which are often hypertrophic in the rectum and left colon (14). It is believed this nerve fiber hyperplasia is a compensatory mechanism aimed at mitigating the loss of inhibitory nerve signals that normally regulate peristalsis. Noteworthy, these fibers are associated with SCPs, and thus these cells have recently received considerable attention due to their presence in the affected bowel and their possible regenerative potential (15, 16). However, it is unclear how SCPs contribute to ENS diversity and whether these cells have a comparable capacity to form both enteric neurons and glia.

Single cell transcriptomics has accelerated our understanding of the composition of the healthy fetal and adult human ENS, in particular of the neuronal compartment (17–19). Considering that defects in the early establishment of this system manifest themselves in childhood in the form of congenital intestinal neuropathies, a better understanding of the composition of the pediatric ENS is required. Here, we leverage human pediatric single cell transcriptomic data from controls and HSCR-affected individuals, to elaborate on the transcriptional diversity of the ENS in health and disease. We also use a transgenic wildtype and a *ret*^−/−^ zebrafish model to further explore the neurogenic potential of SCPs as a novel therapeutic option for HSCR.

## Results

### Single cell RNA sequencing reveals enteric glia heterogeneity of the pediatric human intestine

Full-thickness ileum and colon tissue samples were collected from seven children without an intestinal motility disorder (Fig 1A, Supplemental Table 1). Since ENS cells are a rare population within the intestine (< 2%), an enrichment strategy to isolate and use these cells as input for scRNAseq was developed based on CD56/CD90 expression (Fig 1B) (20). This strategy yielded 33,692 sequenced pediatric ENS cells, of which 456 were neurons and 33,289 were enteric glia. Enteric glia were broadly grouped into two main classes containing respectively two and six populations. The first class, exhibited a gene signature indicative of canonical enteric glia, based on expression of markers related to neuromuscular transmission, such as *ENTPD2,* as well as *APOE* which has been linked to enteric glia (21). The second group, expressed known Schwann cell markers such as *MAL*, *MPZ* and GFRA3 (22–24). We thus termed these two main classes canonical enteric glia 1-2 and Schwann-like enteric glia 1-6, respectively (Fig 1C-1D). Ileum and colonic enteric glia cells showed comparable cluster heterogeneity (Supplemental Fig 1) and thus, all downstream analyses were performed on the integrated ileum and colon data.

**Figure 1.**
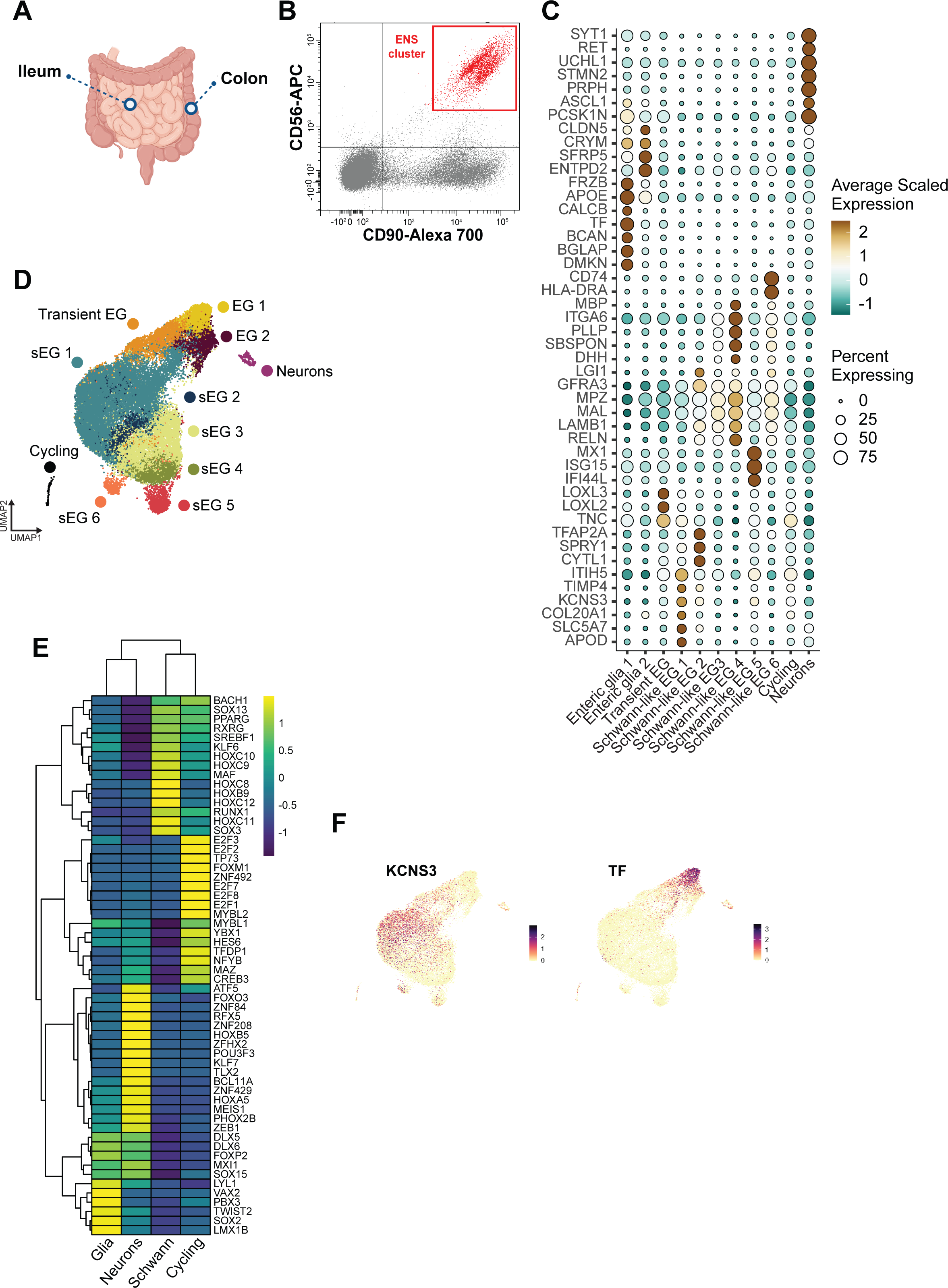
A single-cell transcriptomic landscape of the pediatric Enteric Nervous System. **A)** Schematic of human gut tissue sampling. Biopsies were obtained from ileostomy and colostomy reversals. **B)** ENS cells were enriched via CD56^+^CD90^+^ in a Fluorescence Activated Cell Sorting (FACS) plot. **C)** Dot plot showing expression of differentially expressed genes across cell clusters. **D)** UMAP visualization showed unsupervised clustering of healthy control ileum and colon ENS cells. Indicating the canonical enteric glia clusters 1-2 (EG 1-2) and the Schwann-like enteric glia clusters 1-6 (sEG 1-6). **E)** Heat map showing the mean of differentially expressed transcription factors stratified by major classes of captured cells. **F)** Feature plots showing the differential expression of *KCNS3* and *TF* in the transient enteric glia (EG) cluster.

To provide more insight into the cellular identity of the two glia classes identified, we performed transcription factor analysis. This uncovered that the two glia classes have mutually exclusive signatures, with expression of various posterior *HOX* genes by Schwann-like enteric glia, whilst canonical enteric glia expressed known markers of ENS development, such as *SOX2* and *PBX3* (Fig 1E) (25, 26). Interestingly, a cluster was identified showing a gradient of transcriptional programs characterized by co-expression of genes enriched in both Schwann-like and canonical enteric glia, such as *KNCS3* and *TF* respectively (Fig 1F). This cluster was thus, considered to be a transitional cell state between canonical and Schwann-like enteric glia termed as Transient enteric glia. Interestingly, this cluster was enriched for markers related to the lysyl oxidase family (*LOXL2* and LOXL3) (Fig 1C), suggesting a role in extracellular matrix organization (27).

### Enteric glia exhibit distinct molecular signatures and can be identified by cluster markers *in situ*

To gain further insights into enteric glia diversity of the two major classes identified, the transcriptional signatures of canonical and Schwann-like enteric glia clusters were characterized. Firstly, we examined the expression of known enteric glia markers (*S100B, PLP1, SOX10 and SOX2)* (Fig 2A). The two major classes could not be distinguished based on these markers. However, canonical enteric glia clusters could be readily identified by the specific expression of *APOE* and *RLBP1* (Fig 2B). Moreover, enteric glia 1 expressed *CALCB* and *TF*, suggesting involvement in calcium signaling and iron transport, respectively (Fig 1C). Enteric glia 2 cells were characterized by expression of markers previously linked to the blood-brain-barrier, such as *GJC3* and *CLDN5* (28), suggesting a role in regulating solute and water concentrations in the immediate neuronal environment (Fig 1C, Fig 2B). Interestingly, a small subset of canonical enteric glia showed enrichment of *ASCL1*, a transcription factor involved in enteric gliogenesis and neurogenesis (29) (Fig 2B). Together, these data suggested that canonical enteric glia expressed genes conferring properties inherent to intraganglionic glia (Type I myenteric plexus (MP) and submucosal plexus (SMP)), such as neuromodulation and neurotrophic support (30). To confirm this hypothesis, we performed RLBP1 immunohistochemistry in pediatric intestinal biopsies collected from terminal ileum. As expected, RLBP1 co-localized exclusively in enteric glia located within the myenteric and submucosal plexi (Fig 2C-D), whilst being absent in enteric glia located at interganglionic connectives (Type II enteric glia) and in the lamina propria (Type III enteric glia).

**Figure 2.**
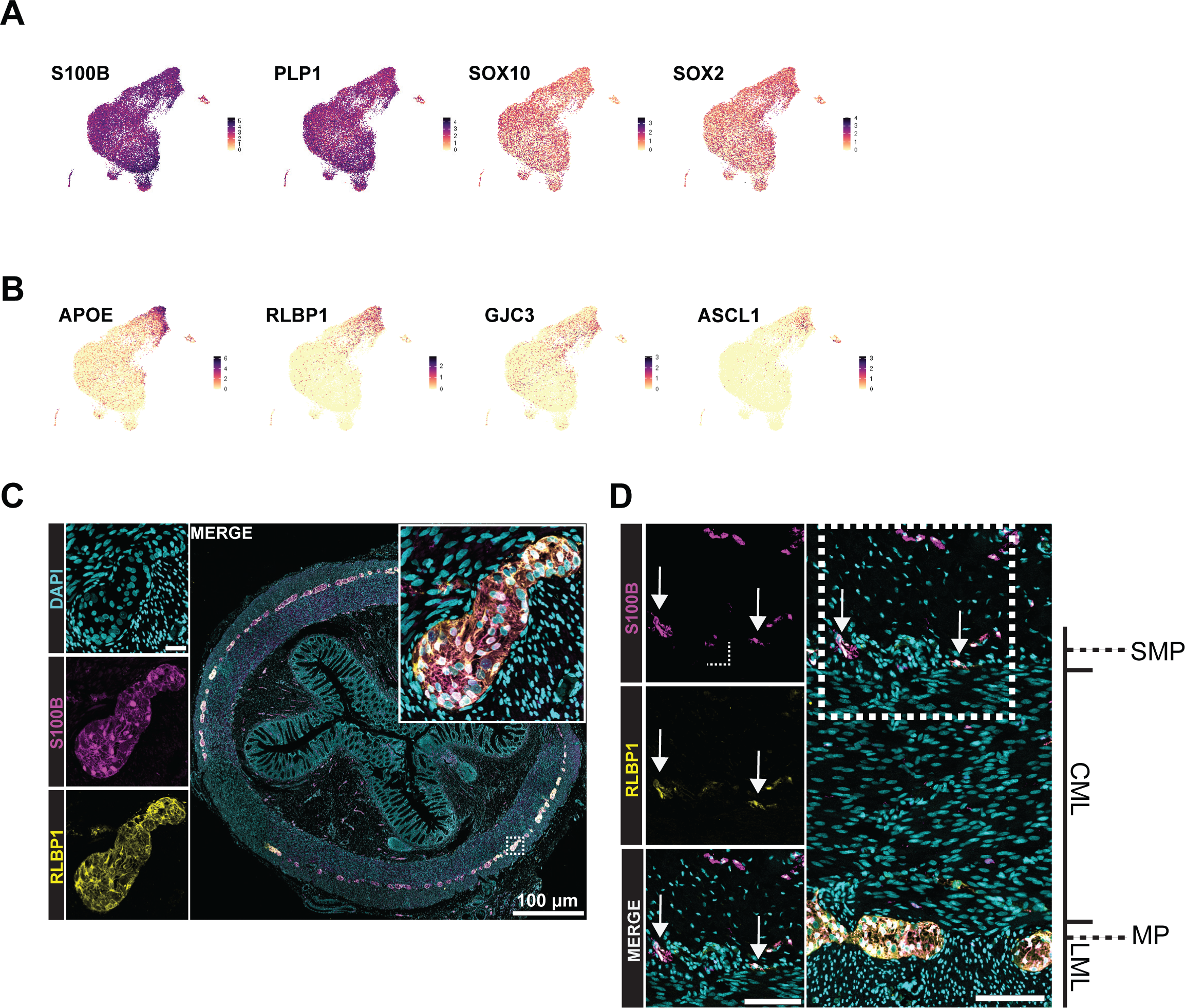
RLBP1 identifies Type I plexus enteric glia. **A)** Feature plots showing differentially expressed genes for the cluster canonical enteric glia 1 (EG 1). **B)** Feature plots showing differentially expressed genes for canonical enteric glia. **C)** Immunohistochemical validation of RLBP1, showing co-localizing to S100B^+^ located in enteric ganglia in the myenteric plexus. **D)** Immunohistochemical validation of RLBP1, showing co-localizing to S100B^+^ located in enteric ganglia in the submucosal plexus. Abbreviation used: MP = myenteric plexus, CML = circular muscle layer, LML = longitudinal muscle layer, SMP = submucosal plexus.

The Schwann-like enteric glia were clustered in six subgroups, Schwann-like enteric glia clusters 1-6. A module score calculated at single-cell level using a publicly available Schwann cell gene list (22), confirmed this similarity (Fig 3A). We found that Schwann-like enteric glia cluster 1 expressed genes involved in ion homeostasis (*KCNC2* and *KCNS3,* Fig 3B), as well as markers reported in peri-synaptic Schwann cells (e.g., *COL20A1* and *APOD*) (31, 32) (Fig 1C, Fig 3B). Interestingly, *SLC5A7*, which encodes for a choline uptake transporter, and has thus far only been described as a marker of cholinergic enteric neurons (17), was selectively expressed in Schwann-like enteric glia cluster 1 (Fig 3B). We exploited this marker to spatially map these cells in the intestine using RNA fluorescent in situ hybridization (FISH), and observed that *SLC5A7*^+^ glia were exclusively located in the muscularis propria, co-localizing with S100B positive cells. Interestingly, Schwann-like enteric glia 1 were characterized by an elongated shape, with fine unbranched processes matching Type IV intramuscular glia (Fig 3C).

**Figure 3.**
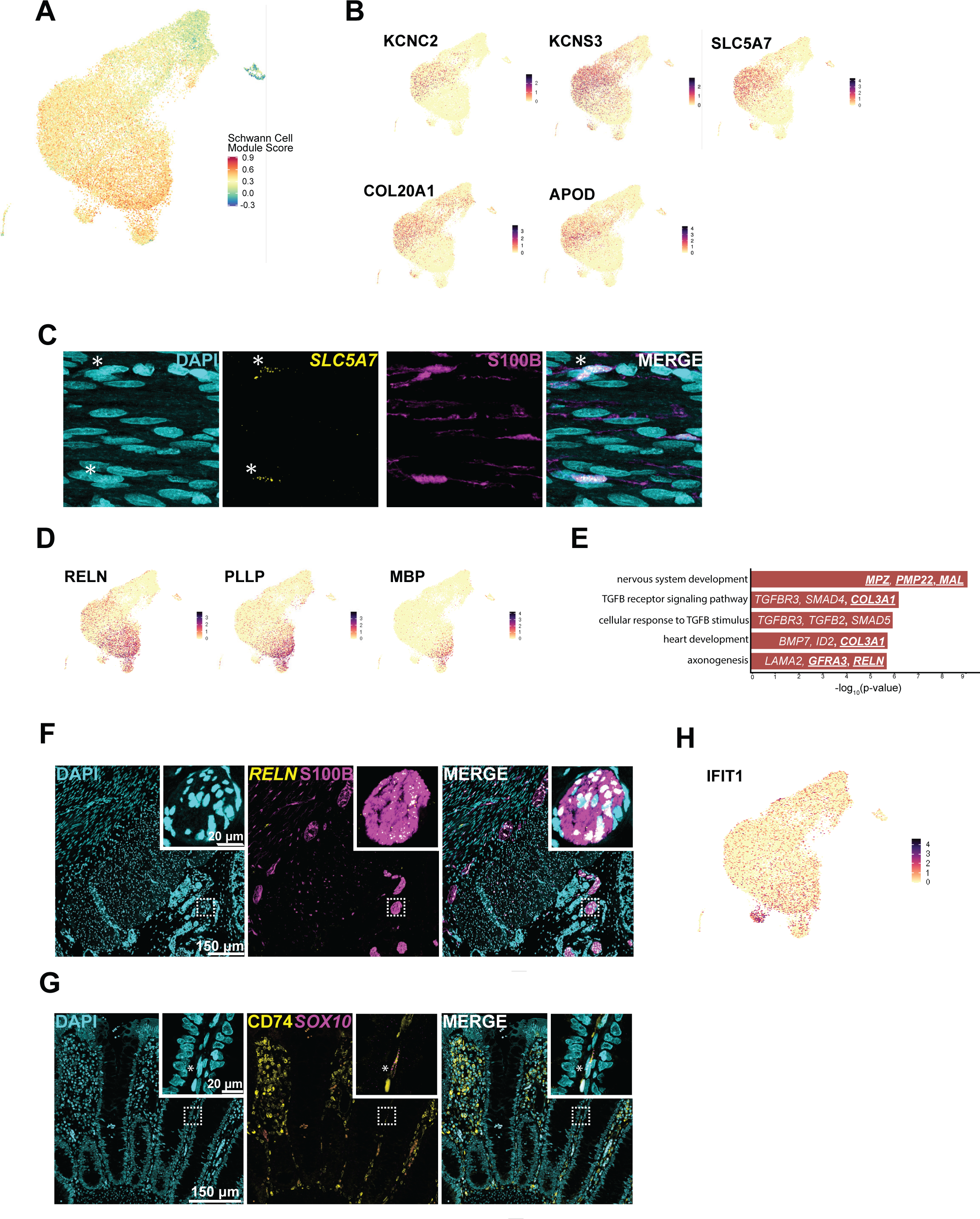
Schwann-like glia represent a heterogeneous type of glia present at various locations along the gut wall axis. **A)** UMAP visualization showing a computed Schwann module score for all cells in our dataset. Schwann-like enteric glia clusters 1-6 have the highest transcriptional resemblance to Schwann cells. **B)** Feature plots showing differentially expressed genes for Schwann-like cluster 1. **C)** Fluorescence in situ hybridization (FISH) combined with immunohistochemistry for S100B, showed *SLC5A7* expression in bipolar Type IV enteric glia. **D)** Feature plots showing the expression of differentially expressed genes for Schwann like enteric glia clusters 3-4. E) Gene ontology analysis for Schwann like enteric glia clusters 3-4 showing enriched biological processes. **F)** FISH combined with immunohistochemistry for S100B, localizes *RELN* expression in extrinsic nerve fibers as well as in enteric glia surrounding blood vessels. G) FISH combined with immunohistochemistry localizes CD74^+^*SOX10*^+^ expression in the lamina propria adjacent to the epithelium. **H)** Feature plots showing the differential expression of *IFIT1* in Schwann-like enteric glia cluster 6.

Schwann-like enteric glia cluster 2 was enriched for *TFAP2A*, suggesting that these cells belong to a precursor state (33, 34) (Fig 1C), whilst Schwann-like enteric glia cluster 3 and Schwann-like enteric glia cluster 4 constituted a transcriptional continuum on which they progressively express differentiation markers (e.g., *RELN*, *PLLP* and *MBP*) (35) (Fig 3D). Gene ontology analysis of these markers, revealed enrichment of processes such as nervous system development and axonogenesis (Fig 3E). FISH showed that *RELN*^+^ glia predominantly localized to extrinsic nerve fibers within the mesentery, but also to glia following along submucosal nerve fibers, as well as around blood vessels (Fig 3F, Supplemental Fig 2). These data suggest that Schwann-like enteric glia cluster 3 and Schwann-like enteric glia cluster 4 correspond to Type III plexus glia (MP/SMP).

Schwann-like enteric glia cluster 5 and Schwann-like enteric glia cluster 6 were the rarest subtypes in our dataset. Schwann-like enteric glia cluster 5 was characterized by a molecular signature indicative of the antigen presentation machinery (e.g., *CD74* and various members of the Major Histocompatibility complex) (Fig 1C). FISH and IHC of *CD74*^+^SOX10^+^ cells revealed their occasional presence in the lamina propria of the mucosa juxtaposed to intestinal epithelial cells (Fig 3G), suggesting that Schwann-like enteric glia cluster 5 contribute to the Type III mucosal glial niche (Type III mucosa). Schwann-like enteric glia cluster 6 was enriched for markers associated with the type I interferon signaling pathway (e.g., *ISG15* and *IFIT1*) (Fig 1C, Fig 3H). However, a selective marker for spatial mapping of this cluster proved difficult to find, as the vast majority of cells in our dataset expressed the same markers as this cluster, albeit at lower levels. Interestingly, a similar interferon related gene signature was recently reported in mucosal enteric glia obtained from ulcerative colitis individuals, indicating that this cluster might constitute a glial cell state induced upon inflammation (36–38). Taken together, these findings indicate that the heterogeneity found in the enteric glial transcriptomic landscape reflects the morphological, topographical and functional diversity of enteric glial cells.

### The proportions of canonical vs. Schwann-like enteric glia are inverted during fetal to pediatric development

Several studies have reported on the developmental dynamics of enteric neurons from fetal stages into later stages of life (39–42). However, there is a paucity of data on the progressive fetal-to-postnatal transition of glial cells. To investigate enteric glia development during these stages, our dataset was integrated with human fetal (6-22 post-conceptual weeks (PCW)) enteric glia data available (18, 19). Labels were assigned based on our pediatric annotations (Fig 4A). The majority of enteric glia present during fetal development were classified as canonical enteric glia 1-2 (Fig 4A-B), and they arose at the earliest timepoints (6-10 PCW) (Fig 4C). In contrast, Schwann-like enteric glia, were less abundant in fetal stages and became the predominant glial class in postnatal samples (Fig 4C). Not surprisingly, the emergence of Schwann-like enteric glia cluster 5-6, which was linked to antigen presentation and immune modulation, only became apparent in our postnatal dataset.

**Figure 4.**
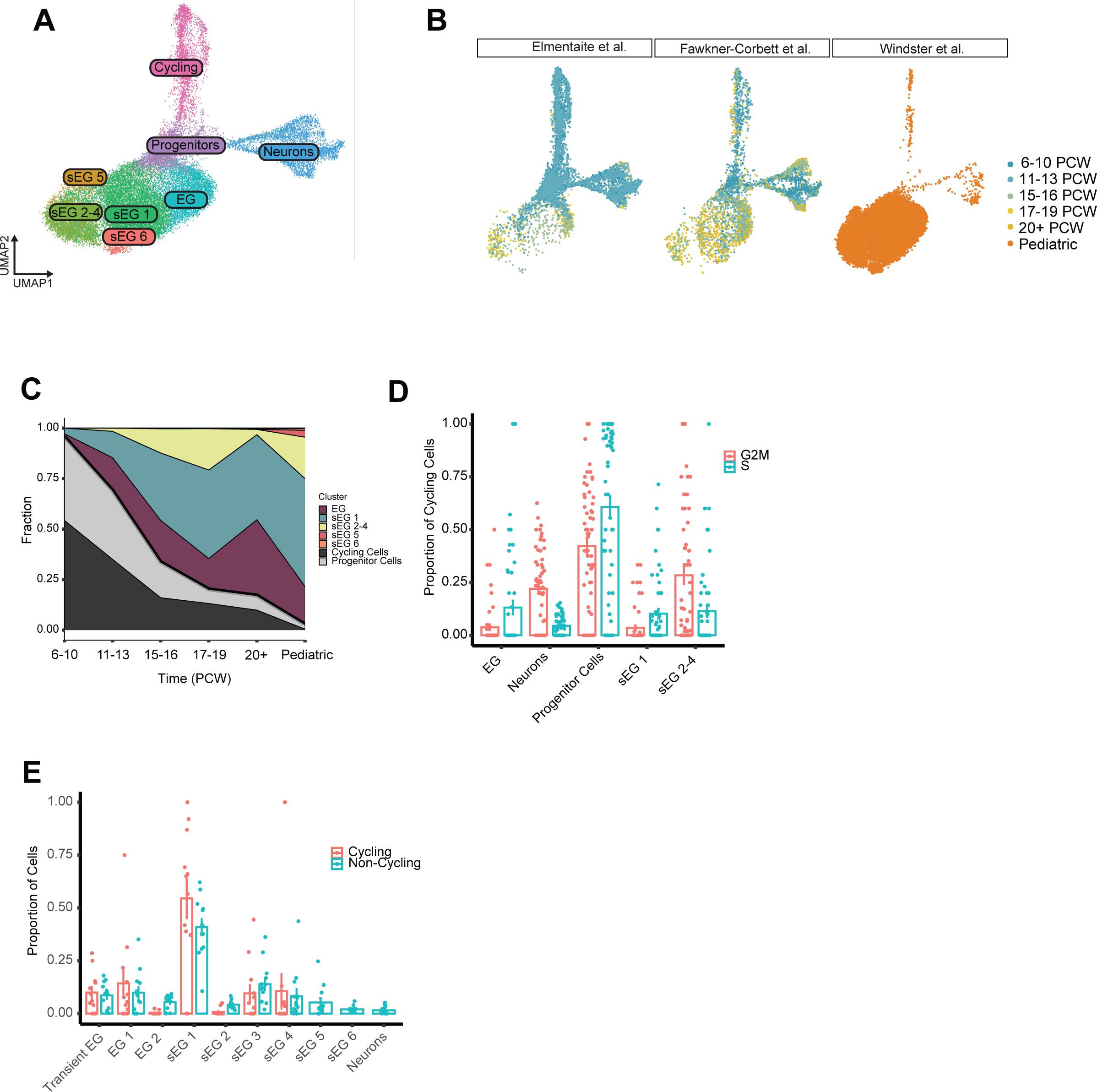
Fetal to pediatric development of the human ENS. **A)** UMAP visualization showing an integration of pediatric and fetal (Elmentaite et al., 2021, Fawkner-Corbett 2021) ENS in which fetal data was cross labeled based on pediatric cluster annotations. **B)** UMAP visualization of each individual data set colored by age groups. Canonical enteric glia (EG) and Schwann-like enteric glia (sEG) develop at later time points compared to neurons. Schwann-like enteric glia cluster 5-6 are less abundant in fetal stages. **C)** Proportion plot depicting the relative abundance of glial, cycling and progenitor clusters across developmental time points. Canonical enteric glia (EG) show a stable transition from fetal to postnatal time points, whereas Schwann-like enteric glia (sEG) become the most abundant subclass during gestation. **D)** Bar plot showing the contribution of fetal ENS clusters to the cycling and non-cycling pool. Progenitor cells are the most highly cycling cells, whilst neurons are characterized by exit of the cell cycle phase. **E)** Bar plot showing the contribution of pediatric ENS clusters to the cycling and non-cycling pool. In contrast to fetal stages, Schwann-like enteric glia cluster 1 becomes the main contributor to cycling cells postnatally.

Lastly, the fetal data were used to determine to which extent each cluster contributed to the cycling population, as a proxy for putative glio- and neurogenesis. As expected, in the fetal dataset the progenitor cells showed the highest contribution to cycling cells (Fig 4D). Postnatally however, although the number of cycling cells were limited, Schwann-like enteric glia cluster 1 showed the largest proportion of G2M/S phase cycling cells, suggesting a role in replenishment of the enteric neuroglial pool for these cells in the postnatal gut (Fig 4E).

### Schwann-like enteric glia are selectively preserved in aganglionic colon of individuals with HSCR

Having established a normal human pediatric enteric glia atlas, we investigated how these cell populations are affected in the context of disease. Full-thickness biopsies of aganglionic and ganglionic regions, but not transition zone, from three HSCR individuals were collected following pull-through surgery, and were analyzed by scRNA-seq (Supplemental Table 1). The single cell transcriptome obtained for the ganglionic and aganglionic regions were compared to our annotated healthy control dataset. Controls and HSCR derived material were not matched with regards to colonic location. Strikingly, whilst the ganglionic colon contained all enteric glia subsets and neurons found in healthy controls, the aganglionic colon was devoid of Canonical enteric glia 1 and 2, transient enteric glia and neurons. Only Schwann-like enteric glia cluster 1-6 were conserved in the aganglionic region (Fig 5A). As aganglionic bowel is devoid of ganglia, and thus should also be lacking ganglia-associated enteric glia, this observation further supports our initial classification of Glia 1-2 as intra-ganglionic glia. FISH for *SLC5A7*^+^ revealed that Schwann-like enteric glia cluster 1 cells appeared to be similarly dispersed between the muscle layers in biopsies from the aganglionic segment of HSCR individuals, as in healthy controls (Fig 3C and 5B). Also, the presence of *RELN*^+^ Schwann-like enteric glia clusters 3 and 4 was validated in the aganglionic segment, and these cells were found localized on hypertrophic nerves (Fig 5C).

**Figure 5.**
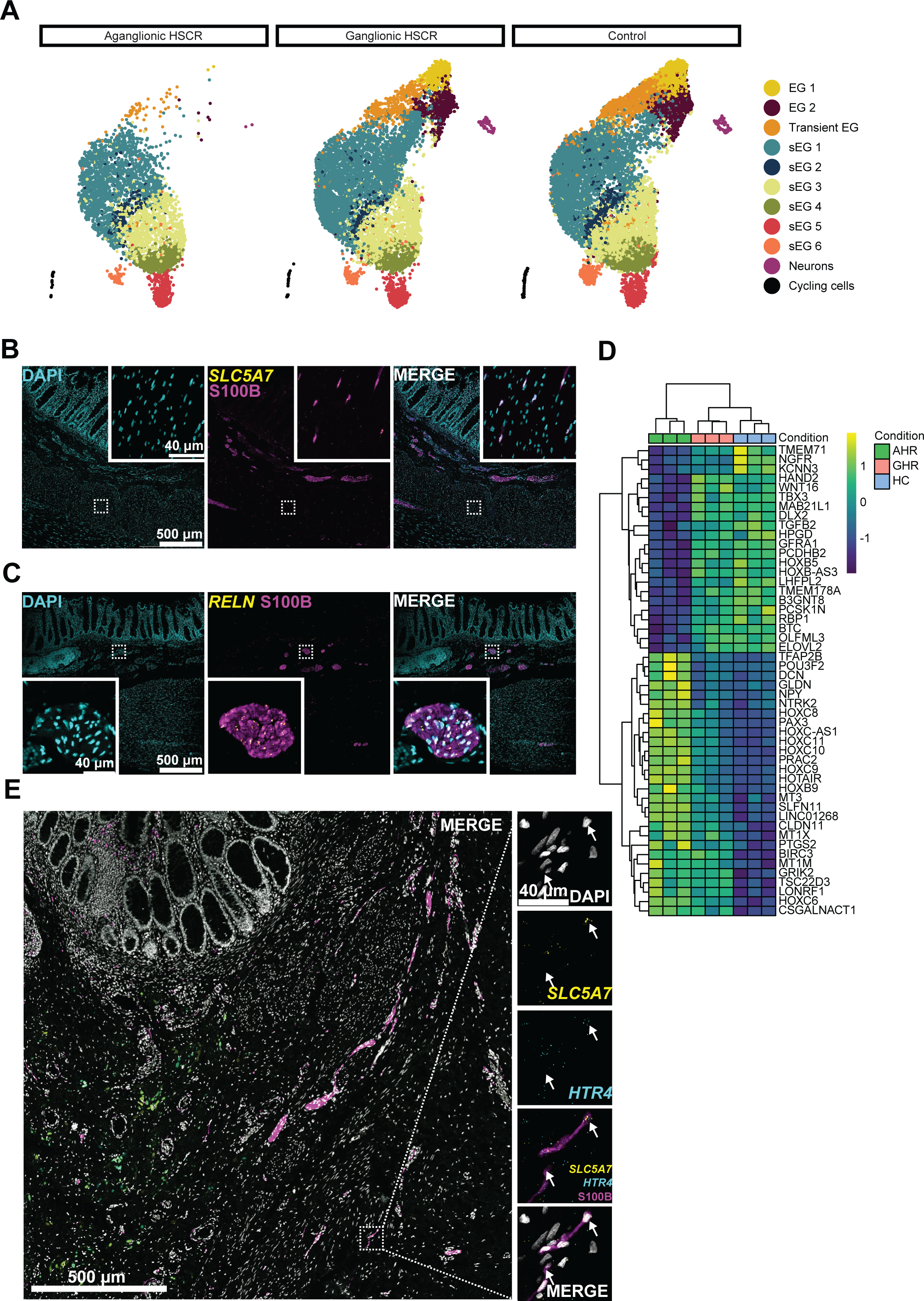
The molecular landscape of the ENS of HSCR individuals is dominated by Schwann-like enteric glia. **A)** UMAP visualizations of aganglionic, ganglionic and control ENS cells. Canonical enteric glia 1-2 and neurons are absent in aganglionic HSCR colon. **B)** Fluorescence in situ hybridization (FISH) combined with immunohistochemistry for S100B localizes *SLC5A7* expression in Schwann-like enteric glia in the aganglionic HSCR colon. **C)** FISH combined with immunohistochemistry for S100B localizes *RELN* expression in glia present in extrinsic hypertrophic nerve fibers in the aganglionic HSCR colon. D) Heat map showing differentially expressed genes stratified by condition. **E)** FISH for *SLC5A7* combined with immunohistochemistry for S100B shows expression of *HTR4* in Schwann-like enteric glia present in the aganglionic colon. Right arrow: S100^+^*RELN^+^SLC5A7^+^* cell. Left arrow: S100B^+^*RELN^-^SLC5A7^-^* cell. Abbreviations: AHR = Aganglionic Hirschsprung Region, GHR = Ganglionic Hirschsprung Region, HC= Healthy Colon

Differential gene expression analysis on Schwann-like enteric glia between aganglionic and healthy control samples, showed reduced expression of signaling and transcription factors such as *GFRA1* and *HAND2* (Fig 5D). However, an increased expression of gene signatures associated with neural development, was found in the aganglionic condition. *PAX3*, a known marker of enteric ganglia formation(43, 44), as well as drivers of neuronal specification, such as *NTRK2* and *CLDN11,* were also enriched in the aganglionic colon (Fig 5D). These findings could suggest a compensatory mechanism at play, in which enteric glia from the aganglionic region attempt to de-differentiate into a more progenitor-like state with the purpose to drive neurogenesis and/or gliogenesis.

Serotonin (5-HT) is a neurotransmitter, known to be involved in neurogenesis and neuronal survival. Collective evidence from animal studies have shown that 5-HT, particularly through the activation of 5HT4 receptors (5HTR4), plays a crucial role in the regulation of enteric neurogenesis (45–48). To explore the potential use of 5-HT in HSCR individuals, we examined the presence of 5HTR4 in aganglionic regions. Based on our scRNA-seq dataset, 5-HTR4 was not expressed in any of the cell clusters identified in both ganglionic and aganglionic colon. Similarly, adult and fetal datasets have failed to demonstrate HTR4 expression in enteric glia on a scRNAseq level ((17–19)). However, it is well known that certain transcripts can remain undetected by scRNAseq (49). Using FISH for *SLC5A7* and 5*HTR4*, and immunostaining for S100B, expression of these markers was observed in human Schwann-like enteric glia present in the aganglionic segment of HSCR individuals (Fig 5E).

### Schwann-like enteric glia are preserved in *ret* ^−/−^ zebrafish

To further explore the therapeutic potential of Schwann-like enteric glia, the zebrafish was used as a model organism. Previous studies have shown that in the intestine of both larvae (5 days post-fertilization (dpf)) and adult zebrafish, enteric glia with neurogenic potential are present (50, 51). In addition, SCPs have also been reported at 5dpf and were shown to play a key role in enteric neurogenesis during larvae development, as well as in case of injury (46, 47). However, very little is known about the presence of enteric glia and SCPs in zebrafish in the absence of an ENS. To investigate this, we made use of the available zebrafish HSCR model, *ret^hu2846/hu2846^* (herein referred to as *ret*-/-), and performed scRNA-seq on 5dpf zebrafish intestines presenting with a phenotype reminiscent of total colonic aganglionosis (52) (Fig 6A). The scRNA-seq dataset obtained, was compared to the one we have previously reported, for wild-type 5dpf zebrafish intestines using the same approach (50). As expected, an overall dramatic loss of ENS cells in the *ret*-/-zebrafish was observed (Fig 6B). IPANs (*nmu, tac3a*) and *phox2bb*-neurons (*sv2a, calb2a*) were almost completely absent, as well as inhibitory motor neurons (*vipb, nos1*), differentiating neurons (*tmsb, syk*), actively proliferating cells (*mi67, pcna*) and cells in a more premature state such as the notch responsive clusters (*her4.2.1, notch3*) and migrating cells (*dcn, wnt11r*) (Fig 6B; Supplemental Fig 3A) (25, 53-56). Also, the canonical enteric glia population expressing *slc2a1b, cx43* and *s100b* we recently described (50), was completely lost in the HSCR model (Fig 6B; Supplemental Fig 3B). However, one particular cluster remained unaffected. Based on the transcriptional expression of *mmp17b, fabp7a, clic6*, *anxa1a* and *col18a1a,* this cluster seems to represent SCPs (Fig 6B; Supplemental Fig 3B) (57–63). Interestingly, expression of *ret* was not detected in this cluster, supporting the notion that its origin is *ret-*independent (Supplemental Fig 3C).

**Figure 6.**
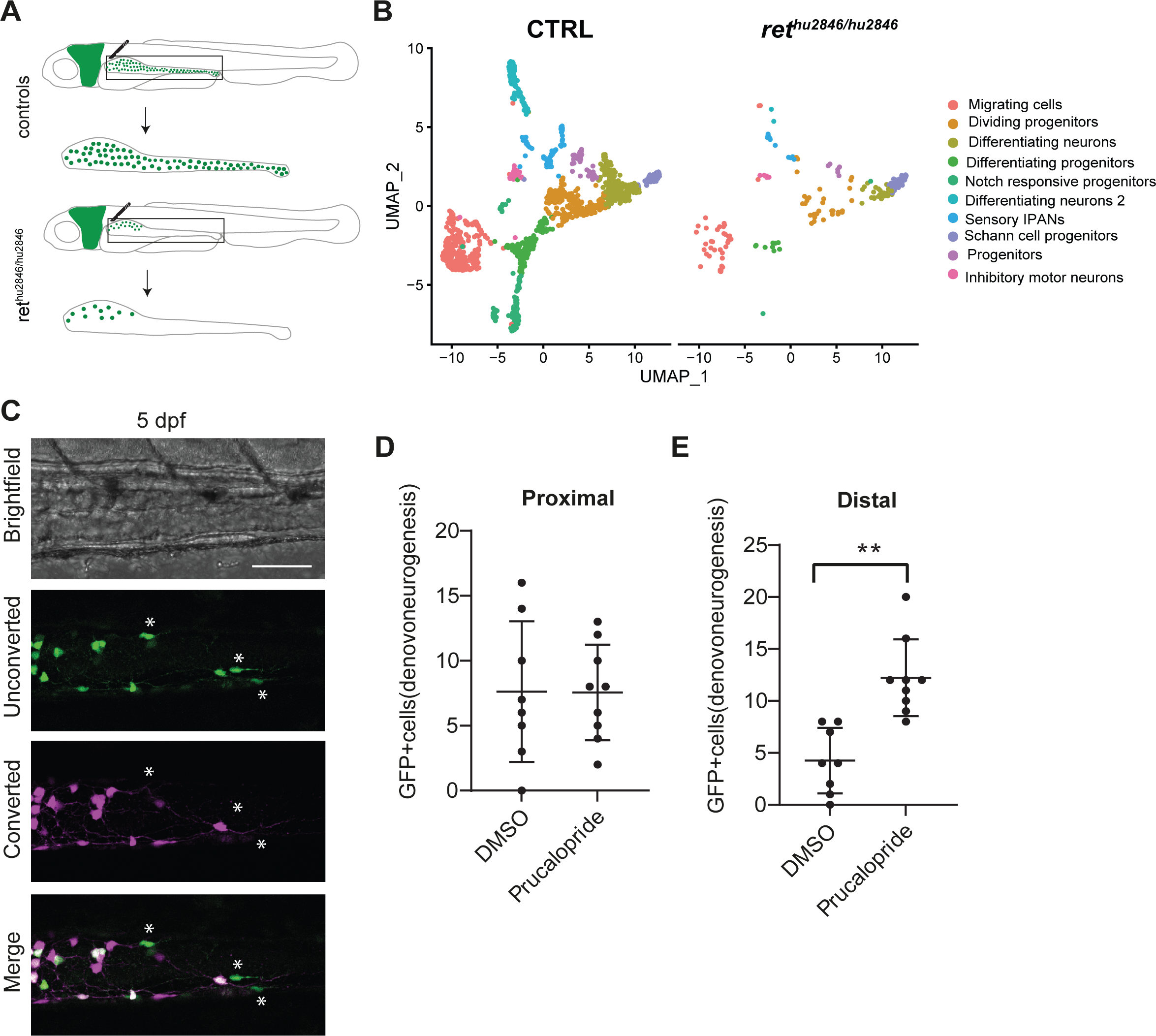
Schwann cell precursor like cells are preserved in *ret*-/-zebrafish and can be stimulated into neurogenesis by prucalopride. **A)** Graphical design of gut isolation for scRNA seq of 5dpf tg(*phox2bb*:GFP) and *ret-/-* tg(*phox2bb*:GFP) zebrafish. **B)** UMAP of ctrl and *ret* mutant ENS subset. **C)** Live-imaging of 5dpf tg(*phox2bb*:kaede) photoconverted zebrafish treated with prucalopride. New *phox2bb*:GFP+ cells are indicated by asterisks (*). **D)** Quantification of *phox2bb*:GFP+ cells in the proximal region of DMSO and prucalopride treated 5 dpf photoconverted tg(*phox2bb*:kaede) zebrafish. **E)** Quantification of *phox2bb* GFP+ cells in the distal region of DMSO and prucalopride treated 5 dpf photoconverted tg(*phox2bb*:kaede) zebrafish.

### Prucalopride promotes neurogenesis in the *ret*^+/-^ fish, partially rescuing the HSCR phenotype

To examine the neurogenic potential of SCPs in the intestine of the HSCR zebrafish, we used a model previously established, in which photoconversion of the tg(8.3*phox2bb:*kaede) transgenic line in the intestine enables detection of newly formed *phox2bb+* neurons (47). With this assay, *phox2bb+* cells present in the intestine are photoconverted (red), allowing identification of new *phox2bb*+ cells derived from different progenitors (green). El-Nachef and Bronner showed that these newly formed cells derive from SCPs (47). Here, we performed photoconversion of all *phox2bb+* cells in the intestine followed by time lapse imaging, using *ret-/-* and *ret+/-* zebrafish presenting respectively, with a phenotype reminiscent of total colonic aganglionosis and short-segment HSCR (Supplemental Fig 4A). The emergence of unconverted *phox2bb*:kaede+ cells was observed in both *ret-/-* and *ret+/-* zebrafish (Supplemental Movie 1; Supplemental Movie 2), showing that *de novo* neurogenesis from SCPs occurs in our HSCR model. Considering that prucalopride, a selective 5-HTR4 agonist, has been shown in zebrafish, to promote enteric neurogenesis from SCPs after injury (47), 4.5 dpf larvae were treated with this compound after photoconversion. Dimethylsulfoxide (DMSO) was used as vehicle control. No increase in *de novo* neurogenesis was observed in the *ret-/-* zebrafish at 5dpf. However, a significant increase in *de novo phox2bb*:kaede+ cells (unconverted; green) was observed in the *ret*+/-model, in the presence of prucalopride (Fig 6C). This effect was specifically evident in the distal intestine (Fig 6D-E; n=8 CTRL, n=9 prucalopride p=0.0067 distal, 0.7746 proximal, 0.0309 total). To study the effects of prucalopride over a longer period of time, *ret+/-* and *ret-/-* tg(*phox2bb*:GFP) zebrafish were treated for three days, from 4 (T=0) to 7 dpf (T=1) (Supplemental Fig 4B). While there was no stimulation of neurogenesis observed in the *ret-/-* zebrafish, a significant increase in the number of *phox2bb+* enteric neurons per 100 µm, was observed in treated *ret^+/-^* zebrafish (short-segment) at 7dpf, in comparison to untreated fish (n=23 DMSO, n=21 prucalopride, p=0.0495 (Supplemental Fig 4C).

## Discussion

Using the power of scRNA-seq we describe for the first time the diverse enteric glia pool in the human pediatric intestine and show that two major enteric glia classes are present. The first class composed by enteric glia 1-2 seems to represent the canonical and traditionally, the most studied type of enteric glia. The second class identified as Schwann-like enteric glia, present with a transcriptome akin to Schwann cells, and were classified as Schwann-like enteric glia 1-6. Based on our results, canonical and Schwann-like enteric glia can be distinguished based on their transcriptional profile, morphology and location: enteric glia 1 and 2 are solely intra-ganglionic, while Schwann-like enteric glia clusters 1-6 are located both intra and extra-ganglionic, such as in the intramuscular and mucosal layers. Interestingly, Schwann-like enteric glia are present in relative low numbers during fetal development, but they become the predominant class in the pediatric intestine postnatally. Although we cannot exclude that these differences can be related to our enrichment method (20), our results are in line with what has been previously described in mice (6), especially because we also showed that Schwann-like enteric glia express 5-HTR4, which was reported to be instrumental for survival/neurogenesis of the adult ENS in vivo (46). Moreover, the Schwann-like enteric glia clusters 5 and 6 detected in the pediatric but not in the fetal intestine, show a transcriptome involved in antigen presentation and immune modulation, which support previous studies stating that enteric glia diversification is contingent on the postnatal maturation of the gut microbiota and immune system (38, 64-66). Taken together, our data suggest that enteric glia ontogeny seems to change after birth and that Schwann-like enteric glia are the major contributor to glia diversity in the healthy pediatric intestine.

The presence of hypertrophic extrinsic nerve fibers in the aganglionic segment, is a known feature of HSCR (14). They are concentrated in the rectum and contain many Schwann cells which have recently been shown to contribute, at least in part, to gliogenesis and neurogenesis in several HSCR mouse models (6, 16). Intriguingly, SCPs derived from hypertrophic nerve bundles of HSCR individuals could be expanded *in vitro* and transplanted into recipient mice where the ENS was ablated using diphtheria toxin (15). These cells were even able to differentiate into glial and neuronal phenotypes (15). Here, we provide transcriptional evidence that Schwann-like enteric glia are indeed preserved in the absence of enteric ganglia, and show suggestive results of the neurogenic capacity of these cells. However, when comparing the transcriptomic profiles of the Schwann-like enteric glia clusters identified in the pediatric human intestine with the SCPs present in zebrafish, it became evident that the different developmental stages led to distinct profiles, with the zebrafish showing a more precursor-like identity, as expected.

In the last couple of years, there has been a search for compounds that could stimulate ENS progenitor cells. For example, treatment of embryonic *EdnrB^NCC−/−^* mouse colon with a laminin-β1 analog (YIGSR), reversed the aganglionic phenotype normally present in these animals (67). Moreover, GDNF was found to induce enteric neurogenesis in the *HolTg/Tg;Dhh-CreTg/+;R26YFP/+* mouse model for trisomy 21-associated HSCR (16). A similar effect was also observed in human *ex vivo* cultures of intestinal tissues collected from HSCR individuals (16). However, the efficiency of GDNF was described to be highly variable and attempts to rescue lethality in the *Ret9/−* mouse model failed. Here, we show that prucalopride, a selective serotonin receptor agonist known to target SCPs (47), is able to promote enteric neurogenesis in the *ret+*/-short segment HSCR zebrafish, providing the first evidence of a partial rescue of the HSCR phenotype in a *ret* genetic model. Neurogenesis was most prominent stimulated in the distal part of the gut, which is in line with a recent report stating an inverse correlation between Schwann-derived neurogenesis and neuronal density in the intestine (5). This is also in line with the fact that the neurogenic potential of SCPs, may differ at different locations in the colon, and is known to be higher in the aganglionic rectum than in aganglionic transverse colon (68). We were however, unable to induce neurogenesis in the *ret-/-* total colonic HSCR zebrafish. This result was not unexpected, as it supports the notion that *RET* expression is essential for neuronal differentiation at later stages. Moreover, it has been suggested that the presence of at least some intrinsic neurons is important for Schwann-derived neurogenesis to take place (6). Considering that the majority of HSCR individuals are heterozygous for *RET* pathogenic variants and are characterized by short segment aganglionosis (80%) (13), we believe that prucalopride can have a positive impact in the treatment of HSCR. Since expression of 5-HTR4 was detected in *SLC5A7*+ Schwann-like enteric glia present in the aganglionic bowel of HSCR individuals, and prucalopride has been approved by the European Medicine Agency (EMA) for the treatment of chronic intestinal pseudo-obstruction (CIPO) (69), its potential implementation could even be accelerated. However, future clinical studies are required to evaluate the use of prucalopride as a new therapeutic option for HSCR.

## Acknowledgements

We thank R.M. Hoogenboezem (Department of Hematology, Erasmus MC Cancer Institute) for processing our raw single-cell data and the Optical Imaging Center (OIC) of the Erasmus MC for maintenance and assistance with the use of confocal microscopes. We would also like to thank R.P. Kapur (Department of Pathology, University of Seatle), W. El-Nachef, M.P. Verhagen (Department of Pathology, Erasmus Medical Center), O.J.M. Schäffers (Department Reproduction and Development) and A.J. Burns (Takeda) for insightful discussions. TSB was supported by the Netherlands Organisation for Scientific Research (ZonMw Vidi, grant 09150172110002), an Erasmus MC Fellowship 2017, and Erasmus MC Human Disease Model Award 2018. VM was supported by the Netherlands Organisation for Scientific Research (ZonMw Vidi, grant). NK and MMA were supported by the Sophia Foundation (S17-18; S20-63; S22-76). Funding bodies did not have any influence on study design, results and data interpretation or final manuscript.

## Data availability

All single cell data reported in this work will be deposited in the Gene Expression Omnibus (GEO) database.

## Methods

### Human material

Human intestinal tissue was collected from resection specimens of children (1 month to 1 year of age) who underwent surgery for ileostoma or colostoma reversal, due to necrotizing enterocolitis or anorectal malformations (control group, n=7). Sampling was performed from macroscopically healthy bowel tissue, following ileostomy and colostomy closure. Supplemental Table 1 describes the diagnosis and major characteristics of all donors included in this study. HSCR individuals admitted at the Pediatric Surgery Department of the Erasmus University Medical Center – Sophia Children’s Hospital were also included (n=3). Biopsies of approximately 2-3 cm^2^ were taken from discarded tissue of resection specimens (ganglionic and aganglionic regions) and rinsed in ice cold PBS. Intestinal tissue was then minced thoroughly with a scalpel, into small pieces of approximately 5 mm^2^, and placed in a cryovial containing 1.5 ml CryoStor cell cryopreservation media (C2874, Sigma). Vials were stored at −80°C for later use.

### Tissue dissociation

Cryopreserved tissue was rapidly thawed in a 37°C water bath for approximately 3 minutes, and subsequently washed with ice cold PBS. Tissue dissociation was performed as described before (20). In short small pieces of full-thickness specimens were placed in a gentleMACS C-tube (130-093-237, Miltenyi Biotec) in digestion solution and dissociated in the gentleMACS Octo Dissociator (130-095-937, Miltenyi Biotec) for 1 hour (program 37C_h_TDK_1). The tissue was thereafter gently triturated with a needle and syringe and subsequently strained through a 70 µm cell strainer (130-098-458, Miltenyi Biotec).

### Enrichment of ENS cells from total gut

A fluorescence activated cell sorting (FACS) approach was used to enrich for enteric neurons and glia, as described before (20). In short, viable CD56^+^/CD90^+^/CD24+ cells were isolated from a heterogeneous single cell suspension obtained from tissue dissociation.

### Droplet-based scRNA-seq

Following FACS, human and zebrafish samples underwent droplet based scRNAseq using the 10x Chromium single cell platform RNA Libraries were prepared using the Chromium Single Cell 3’ Reagents kits (10x Genomics) as described by the manufacturer according to their user guide. Quality of the libraries was determined using an Agilent High Sensitivity DNA kit (Agilent Technologies). Finalized libraries were sequenced on a Novaseq6000 sequencing platform (Illumina) for 28-10-10-90 cycles, targeting a minimal read depth of 20000 reads/cell per sample.

### Processing FASTQ reads into gene expression matrices

Cellranger software (version 6.0.1) from 10X Genomics was used to process, align and summarize unique molecular identifier (UMI) counts against hg38 (10x reference: refdata-gex-GRCh38-2020-A) human reference genome. Matched hashing antibody data were processed together with scRNA-Seq using feature barcoding workflow, with oligonucleotide tagged sequences provided in **Supplemental Table 2**. Raw UMI count matrices from Cellranger outputs were imported into R for further processing. For each sample, cell calling was performed using ‘emptyDrops’ (68) function from DropletUtils on the full raw count matrices of all barcodes in the 10x barcode whitelist to distinguish cells from empty droplets containing only ambient RNA. Droplet barcodes for which a high percentage of total UMIs originated from mitochondrial RNAs were filtered out, as well as low total UMI count barcodes. Seurat R package (69) was used for further analysis.

### Hashing

Hashing antibody UMI count matrices were filtered to keep only 10x cellular barcodes from cells passing QC and filtered antibody matrices were used to demultiplex samples as described in Stoeckius et al. (70). Briefly, counts were normalized using centered log ratio transformation and an initial clustering solution was obtained using clara k-mediods clustering with k = 1 + number of samples in the pool. A negative binomial distribution was fit for each hashing antibody. A positive threshold was defined at 99th percentile for each antibody, with cells above this threshold considered positive. Cell sample-of-origin was assigned for each cell based on individual hashtag thresholds. Multiplets were defined as cells positive for multiple antibodies and filtered out from further analyses.

Demultiplexed singlets from separate 10x reactions/samples were merged together and expression values were normalized for total UMI counts per cell. Highly variable genes were identified by fitting the mean-variance relationship and dimensionality reduction was performed using principal-component analysis. Scree plots were used to determine the number of principal components to use for clustering analyses for each pool. Cells were then clustered using Louvain algorithm for modularity optimization using kNN graph as input. Cell clusters were visualized using UMAP algorithm (71) with principal components as input and n.neighbors = 30, spread = 1.5 and min.dist = 0.05.

### Batch correction

Following examination of initial clustering solution, Harmony algorithm (72) was used to correct for batch effects. Merged cell clustering and visualization of cells was then performed as before using Louvain and UMAP algorithms, using harmony dimensionality reduction as input instead of principal components. Batch corrected clusters were compared with cell types obtained from individual 10x reactions and uncorrected data, to ensure that cell type heterogeneity was not lost due to batch correction. Batch-corrected clusters were annotated based on canonical marker gene expression with reference to previously published gut scRNA-Seq datasets (37, 73, 74). Contaminant cell populations, corresponding mainly to fibroblasts, were removed from further analysis to retain only glia and neurons. One cluster of actively proliferating cells was identified based on expression of cell cycle markers (e.g., *MKI67*) and contained proliferating cells from multiple lineages. To retain cycling glia and neurons for further analysis, we carried out a label transfer procedure to assign a phenotype to these cells by sub-setting all non-cycling cell clusters and used these data as a reference to classify cycling cells into broad populations. Classification labels, which we further checked against known marker gene expression within each predicted cell group, were then used to separate cycling glia and neurons from contaminating cell types, only retaining the former for further analysis. All glia and neurons were then re-clustered as described above.

### Clustering and differential gene expression

Cluster markers and differential gene expression were identified using negative binomial generalized linear model statistical testing. In each case, confounding sources of variation from gene detection rate and batch effects were included in the model formula as blocking covariates. Benjamini-Hochberg multiple testing correction was used to correct for multiple testing. Genes with FDR < 5% were considered significantly differentially expressed. R package MAST (75) to identify differentially expressed genes which behave in an on-off manner by testing the continuous and discrete components of the Hurdle model. Gene Ontology and pathway enrichment analyses were performed using the clusterProfiler R package (76), with annotation Dbi R package org.Hs.eg.db used to map gene identifiers. Hypergeometric *P* values were adjusted in each case for multiple testing using Benjamini– Hochberg correction as before. The results were visualized using the R packages clusterProfiler and ggplot2.

For fetal meta-analysis, fetal scRNA-Seq data (18, 19) were downloaded from (ArrayExpress: E-MTAB-9533 and GEO: GSE158702) in fastq format and were processed with Cellranger and Seurat as described above to standardise with the dataset generated here. Only colonic and ileal samples were used. Data were integrated using Harmony algorithm as before, correcting for both individual batches and dataset-specific effects. Trajectories on the integrated embedding were fit using Monocle 3 algorithm (77). Seurat objects were converted to Monocle 3 cell_data_set objects and trajectory reconstruction was carried out allowing for closed loops and multiple partitions. The start of the trajectory was denoted as the node within the least differentiated cells enriched for earliest fetal time points and pseudotime was computed along the trajectory from this node.

To detect active transcription factor modules, normalized single-cell gene expression matrix was first filtered to exclude all genes detected in fewer than 20 cells. The RcisTarget database, containing transcription factor motif scores for gene promoters and transcription start sites for the hg38 human reference genome, was downloaded from https://resources.aertslab.org/cistarget/databases/homo_sapiens/hg38/refseq_r80/mc9nr/gene_based/, and the expression matrix was further filtered to include only genes available in the RcisTarget database. The remaining genes were used to compute a gene–gene correlation matrix for co-expression module detection using the random forest-based GENIE3 algorithm (79); the R package SCENIC (78) was used to perform transcription factor network analysis to detect co-expression modules enriched for target genes of each candidate transcription factor from the RcisTarget database. The AUCell package (80) was used to compute a score for each TF module in each individual cell. To identify condition-or cluster specific TF modules, generalised linear models were used to test for condition or cluster dependence of TF AUC values, including batch and gene detection rate as blocking co-variates in the model formula. Resultant p-values were further adjusted for multiple testing using Benjamini-Hochberg multiple testing correction.

### Immunofluorescence, confocal microcopy and image analysis

Immunofluorescence was performed on tissues which were age-matched to our scRNA-seq samples. Formaline fixed paraffin embedded (FFPE) tissue samples were freshly cut at 6 µm thickness and deparaffinized in xylene. Subsequently, sections were rehydrated in a graded ethanol series. Antigen retrieval was performed for 15 minutes at 94-96 °C on a hot plate in 10 mM sodium citrate at pH 6. Primary antibodies were diluted in PBS and sections were incubated for 1.5 hours. Sections were washed in PBS and incubated for 1 hour with secondary antibodies. The slides were then washed and mounted using Vectashield Vibrance Antifade Mounting Medium with DAPI (Vectorlabs, H1800-10). Images were taken on a Leica Stellaris 5 confocal laser scanning microscope.

### RNAscope assay

Formalin Fixed Paraffin Embedded (FFPE) tissue blocks form pediatric intestine age-matched to our scRNA-seq donors, were obtained from the Department of Pathology of the Erasmus Medical Center. Sections of 6 μm-thick were freshly cut and *in situ* hybridization was performed using the RNAscope Multiplex Fluorescent Reagent Kit v2 ((Advanced Cell Diagnostics (ACD), Cat. No. 323100), following the FFPE tissue protocol provided by the kit. To combine immuno-fluorescence with ISH we used the RNA-Protein Co-Detection Ancillary Kit (ACD, Cat. No. 323180) following the manufacturer’s protocol. Images were taken on a Leica Stellaris 5 confocal laser scanning microscope.

### Animal husbandry

The following zebrafish lines were used: transgenic tg(*phox2bb*:GFP)(32), tg(*8.3phox2bb:kaede)* (ref Ian Shepherd), *ret^hu2846/^* ^+^ (ref) and *ret^sa2684/+^* (ref). Zebrafish were kept on a 14/10h light/dark cycle. Embryos and larvae were kept in an incubator at 28.5°C in HEPES-buffered E3 medium. For imaging experiments, fish were treated from 24 hpf onwards with 0.2 mM 1-phenyl 2-thiourea (PTU), to inhibit pigmentation. Animal experiments were approved by the Animal Experimentation Committee of the Erasmus MC, Rotterdam (AVD1010020209425).

### Isolation of zebrafish intestines and pre-processing of intestinal cell suspension for scRNA sequencing

Intestines of 5 (dpf) larvae were isolated and dissociated as described before (50). Cells were then transferred into FACS tubes using a 35 µm cell strainer, and were centrifuged at 700g for 5 minutes at 4°C. The supernatant was removed and the pellets were resuspended in PBS containing 10% FCS. DAPI was added to mark dead cells (1:1000). Live, single cells were sorted into an eppendorf containing PBS with 5% FCS using the FACSAria III sorter (BD Biosciences).

### Single cell RNA sequencing analysis of the zebrafish cells using Seurat

scRNA sequencing analysis was performed using Seurat V3 (69). For analysis, scRNA-seq datasets obtained for the intestines of 5dpf wildtype (50) and *ret-/-zebrafish*, were integrated. Both datasets were down-sampled to 9200 cells to obtain an equal number of cells (nFeature_RNA > 100 & nFeature_RNA <4200 & percent.mito < 0.05), and the Seurat pipeline was used for normalization, integration and downstream analysis. We used 50 dimensions with a resolution of 0.8 for the clustering, and UMAP for the processing of the individual datasets and also for the post-processing of the merged dataset. This led to 49 clusters of which four expressed neuronal and/or enteric progenitor markers (50). These four clusters were selected for a subset analysis, using 20 dimensions with a resolution of 0.5 for the clustering and UMAP. This analysis provided us with 10 clusters, which were again annotated based on the top 10 differential gene expression and literature search.

### Fluorescent imaging and photoconversion of tg(phox2bb:kaede) zebrafish

Imaging was performed as previously described (51). Briefly, zebrafish were sedated using tricaine (0.016%) and placed on an agarose covered petridish. For confocal microscopy, the Leica M165 microscope was used, and larvae were mounted in 1.8% low melting point agarose on the lateral side. All *phox2bb*:kaede+ cells present in the total intestines, were photoconverted using the 405 nanometer (nm) laser as previously described (47). Full photoconversion was confirmed by the sequential scan of the green and red channels with the 488nm and 561nm lasers. For timelapse experiments, full thickness intestines were recorded at two positions per fish every 15 minutes, for a total of 15 1/2 hours.

### Treatment with Prucalopride in zebrafish

For the experiments where the *de novo* neurogenesis was quantified at 5 dpf, larvae were removed from the agarose after photoconversion and kept protected from the light in the 28.5 degrees incubator overnight. Treatment with prucalopride (10µM; MilliporeSigma SML1371) or DMSO was performed, and 16 hours after the same larvae were mounted and imaged again.

To study the effect of prucalopride over a longer time period, *ret*^+/-^ tg(*phox2bb*:GFP) and *ret*^−/−^ tg(*phox2bb*:GFP) zebrafish were treated with 10µM prucalopride from 4 to 7dpf in a 12-wells plate. Fresh prucalopride was added every other day. Controls were exposed to equal volumes of DMSO.

