## Supplementary figures and images for "Human enteric glia diversity in health and disease: new avenues for the treatment of Hirschsprung disease"

### Supplemental Figures

#### Supplemental Figure 1

A

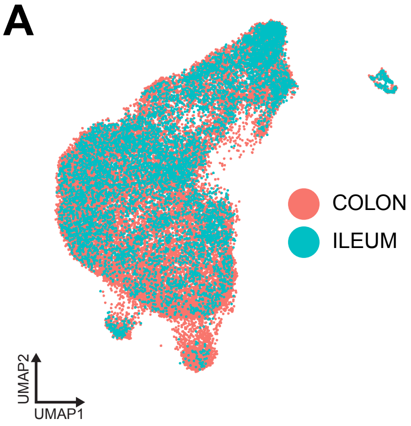


B

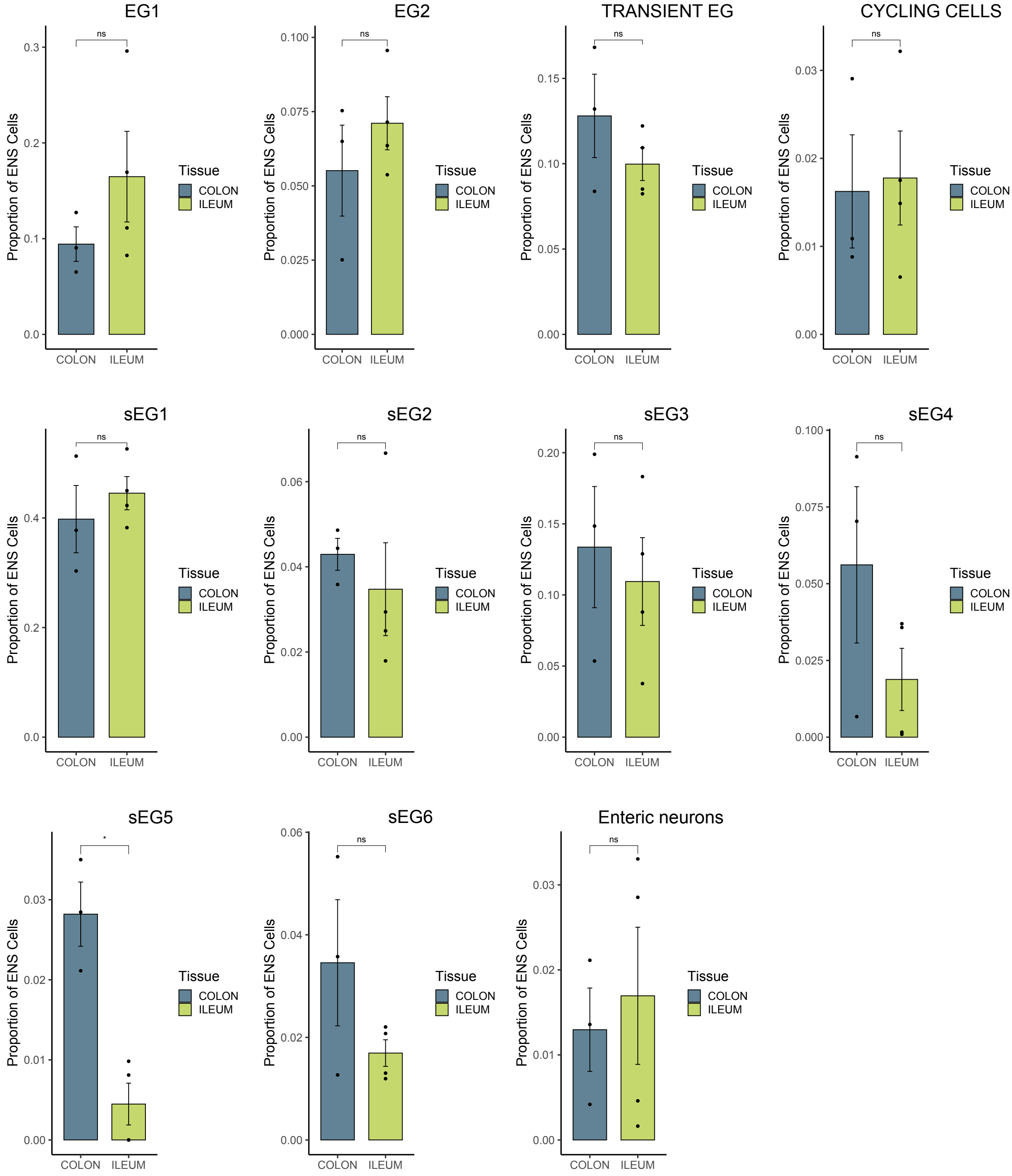

#### Supplemental Figure 2

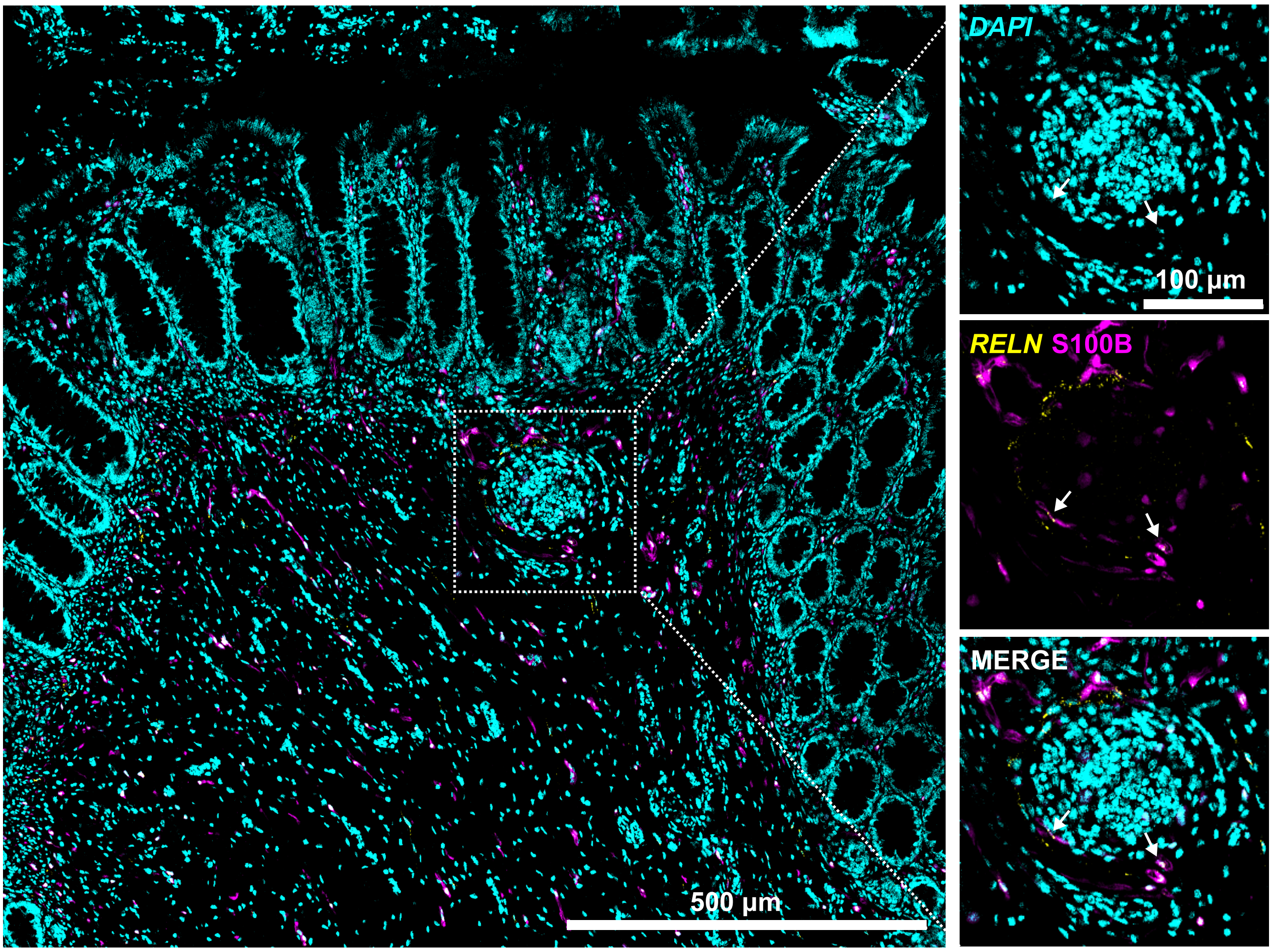

#### Supplemental Figure 3

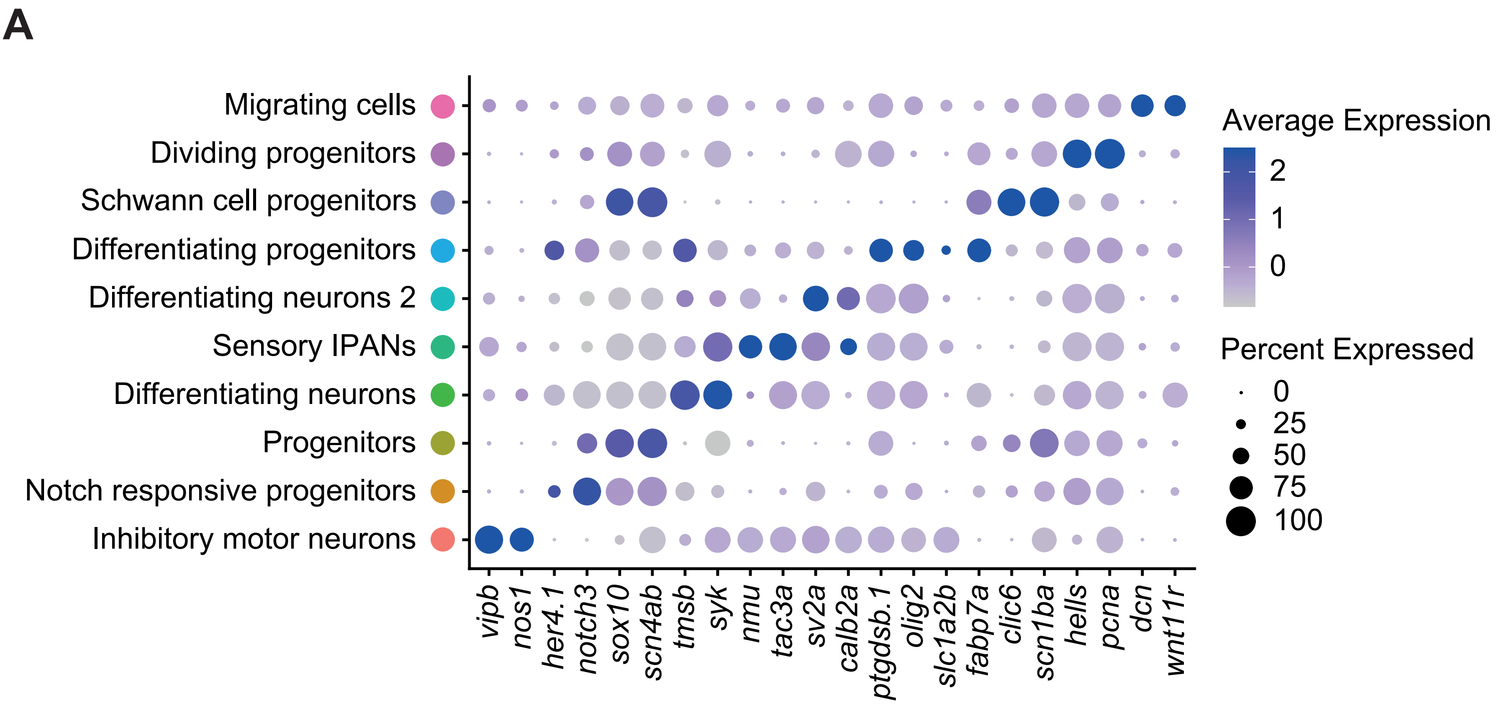


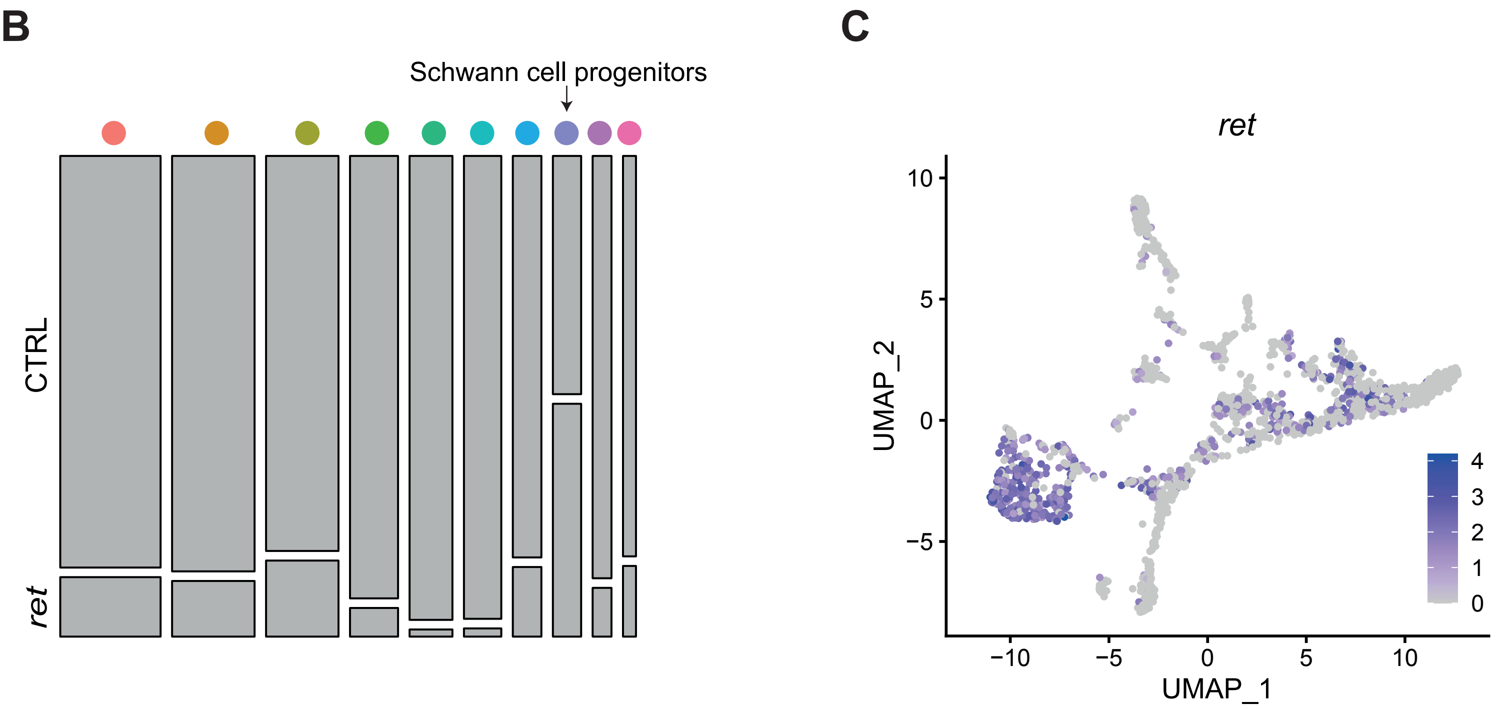

#### Supplemental Figure 4

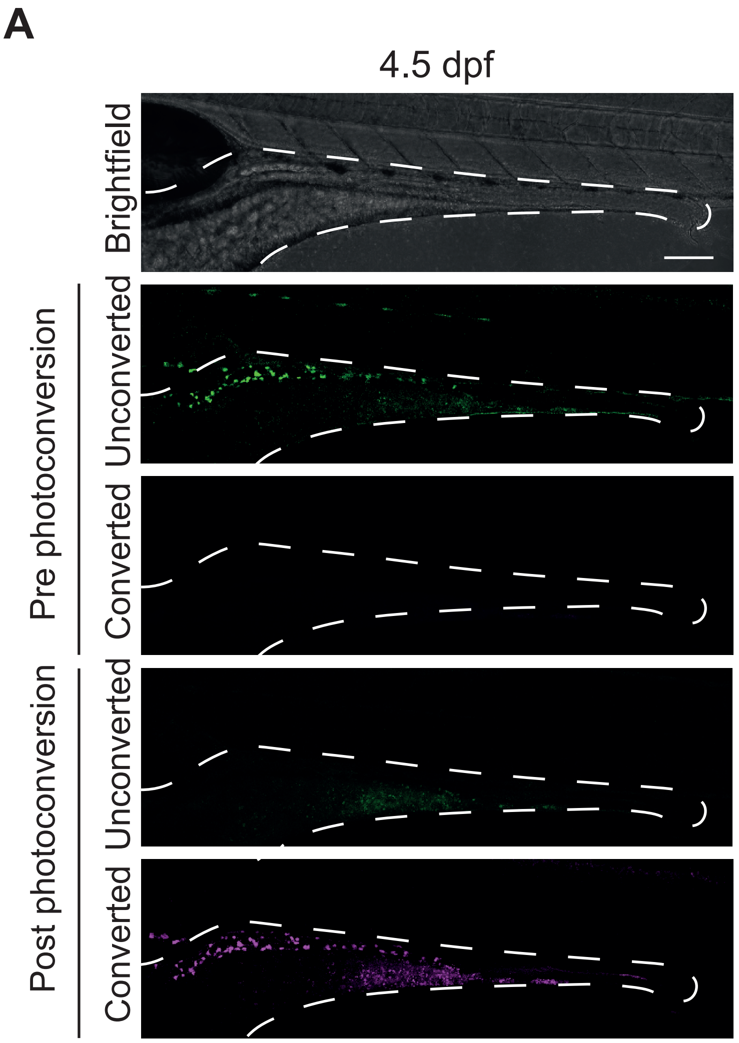


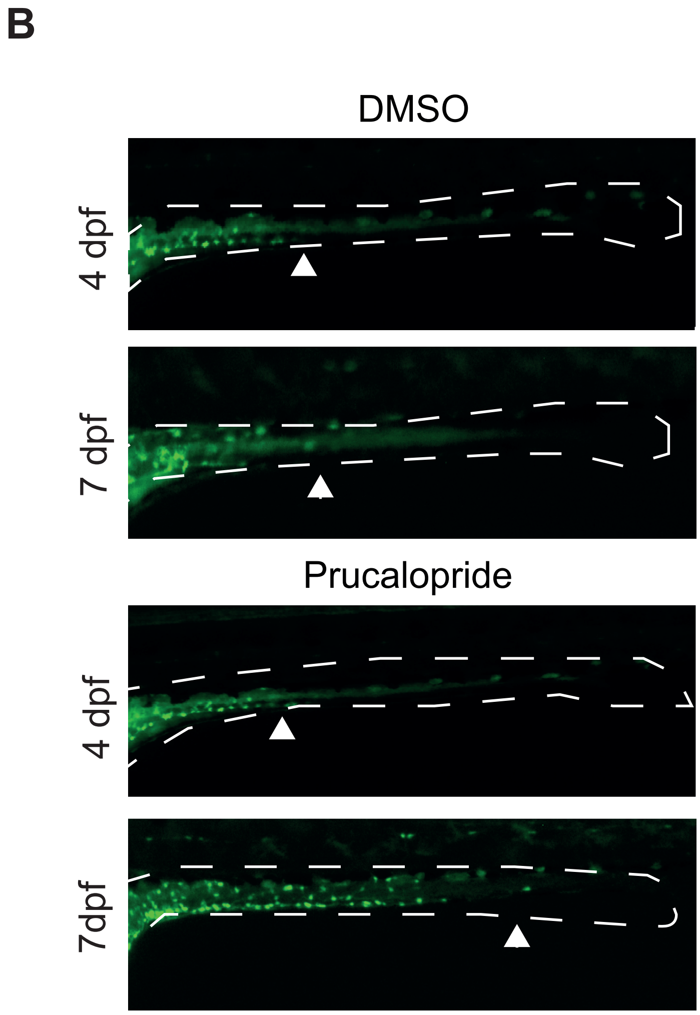


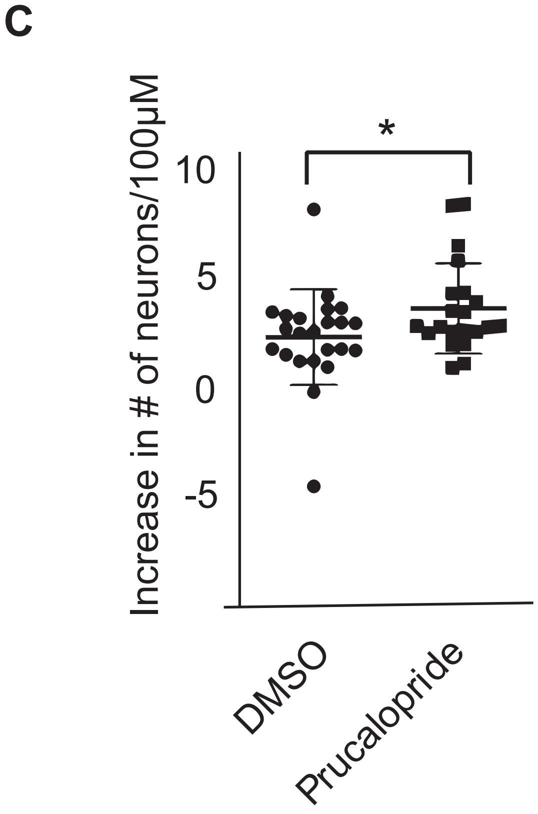
