## Supplemental Table 1 for "Human enteric glia diversity in health and disease: new avenues for the treatment of Hirschsprung disease"

| **Sample ID** | **Sex** | **Age at surgery (months)** | **Indication for surgery** | **Bowel region** | **Macroscopic examination** | **Weight of tissue** | **Cells sorted** | **Cells captured by 10X Chromium Controller** |
| --- | --- | --- | --- | --- | --- | --- | --- | --- |
| Ileum_01 | Male | 3 | Ileostomy closure after necrotizing enterocolitis | ileum | Healthy non-inflamed tissue | ~500mg | 20.000 | 4.608 |
| Ileum_02 | Male | 3 | Ileostomy closure after necrotizing enterocolitis | ileum | Healthy non-inflamed tissue | ~500mg | 216.000 | 7.258 |
| Ileum_03 | Female | 10 | Ileostomy closure after necrotizing enterocolitis | ileum | Healthy non-inflamed tissue | ~500mg | 66.000 | 9.807 |
| Ileum_04 | Male | 2 | Ileostomy closure after necrotizing enterocolitis | ileum | Healthy non-inflamed tissue | ~500mg | 65.000 | 4.777 |
| Colon_01 | Male | 5 | Colostomy closure after anorectal malformation repair | Descending colon | Healthy non-inflamed tissue | ~500mg | 44.000 | 9.922 |
| Colon_02 | Male | 4 | Colostomy closure after anorectal malformation repair | Descending colon | Healthy non-inflamed tissue | ~500mg | 25.700 | 4.579 |
| Colon_03 | Female | 4 | Colostomy colure after necrotizing enterocolitis | Descending colon | Healthy non-inflamed tissue | ~500mg | 10.000 | 1.308 |
| HSCR_01_ganglionic | Male | 4 | HSCR | Rectosigmoid | Macroscopically normal tissue harvested at proximal resection margin at level of ganglionic per-operative frozen section biopsy | ~500mg | 18.100 | 4.908 |
| HSCR_01_aganglionic | Male | 4 | HSCR | Rectosigmoid | Macroscopically thickened tissue at distal resection margin | ~500mg | 13.600 | 3.598 |
| HSCR_02_ganglionic | Male | 2 | HSCR | Rectosigmoid | Macroscopically normal tissue harvested at proximal resection margin at level of ganglionic per-operative frozen section biopsy | ~500mg | 4.100 | 3.301 |
| HSCR_02_aganglionic | Male | 2 | HSCR | Rectosigmoid | Macroscopically thickened tissue at distal resection margin | ~500mg | 20.000 | 1.307 |
| HSCR_03_ganglionic | Female | 24 | HSCR | Rectosigmoid | Macroscopically normal tissue harvested at proximal resection margin at level of ganglionic per-operative frozen section biopsy | ~500mg | 10.000 | 1.898 |
| HSCR_03_aganglionic | Female | 24 | HSCR | Rectosigmoid | Macroscopically thickened tissue at distal resection margin | ~500mg | 7.000 | 1.308 |

**Supplementary Table 1 – Patient diagnosis and major characteristics**
