## Supplemental Table 2 for "Human enteric glia diversity in health and disease: new avenues for the treatment of Hirschsprung disease"

| **Sample ID** | **Hashtag oligo sequence** |
| --- | --- |
| Colon_03 | GTCAACTCTTTAGCG |
| HSCR_02_ganglionic | GTCAACTCTTTAGCG |
| HSCR_02_aganglionic | TGATGGCCTATTGGG |
| HSCR_03_ganglionic | AGTAAGTTCAGCGTA |
| HSCR_03_aganglionic | AGTAAGTTCAGCGTA |

**Supplementary Table 2 – Oligonucleotide tag sequences**
